# Coordinated interplay between palmitoylation, phosphorylation and SUMOylation regulates kainate receptor surface expression

**DOI:** 10.1101/2023.07.24.550331

**Authors:** Busra P. Yucel, Enaam M. Al Momany, Ashley J. Evans, Richard Seager, Kevin A. Wilkinson, Jeremy M. Henley

**Affiliations:** Centre for Synaptic Plasticity, School of Biochemistry, Biomedical Sciences Building, University of Bristol, University Walk, Bristol, BS8 1TD, UK

**Keywords:** kainate receptors, post-translational modification, palmitoylation, phosphorylation, SUMOylation, membrane trafficking

## Abstract

Kainate receptors (KARs) are key regulators of neuronal excitability and synaptic transmission. KAR surface expression is tightly controlled in part by post-translational modifications (PTMs) of the GluK2 subunit. We have shown previously that agonist activation of GluK2-containing KARs leads to phosphorylation of GluK2 at S868, which promotes subsequent SUMOylation at K886 and receptor endocytosis. Furthermore, GluK2 has been shown to be palmitoylated. However, how the interplay between palmitoylation, phosphorylation and SUMOylation orchestrate KAR trafficking remains unclear. Here, we used a library of site-specific GluK2 mutants to investigate the interrelationship between GluK2 PTMs, and their impact on KAR surface expression. We show that GluK2 is basally palmitoylated and that this is decreased by kainate stimulation. Moreover, a non-palmitoylatable GluK2 mutant (C858/C871A) shows enhanced S868 phosphorylation and K886 SUMOylation under basal conditions and is insensitive to KA-induced internalisation. These results indicate that GluK2 palmitoylation contributes to stabilising KAR surface expression and that dynamic depalmitoylation promotes downstream phosphorylation and SUMOylation to mediate activity-dependent KAR endocytosis.

**Significance Statement:** Post-translational modifications (PTMs) are biochemical switches that control substrate protein properties and interactions. In consequence, PTMs are critical regulators of essentially all cellular pathways and are vital for eukaryotic cell survival. In the brain, among other roles, PTMs influence neuronal growth, differentiation, synaptic activity and plasticity. Kainate receptors (KARs) play core roles in all these processes and previous work has shown that the GluK2 subunit of KARs is subject to multiple PTMs. Here, using GluK2 as an exemplar protein, we delineate the sequence, coordination, and consequences of the PTMs palmitoylation, phosphorylation and SUMOylation on KAR surface expression. Our data show how the complex interplay between PTMs dynamically regulates synaptic proteins and neuronal function.

## Introduction

Kainate receptors (KARs) are ionotropic glutamate receptors present throughout the brain at pre-, post-, and extrasynaptic sites, depending on the neuronal subtype and synapse in question. Compared to AMPARs and NMDARs, ionotropic KAR-mediated synaptic responses are highly restricted to subsets of excitatory synapses but, in addition to being ligand-gated ion channels, KARs also signal via metabotropic, G protein-coupled pathways (1–3). Through this combination of signalling modes, KARs play modulatory roles in neurotransmitter release and contribute to postsynaptic depolarisation, and play wider regulatory roles in a range of processes including synaptic plasticity, the formation and maintenance of neural circuits, and excitotoxicity and neuronal cell death (3, 4). Unsurprisingly, therefore, KAR dysregulation and/or dysfunction is strongly implicated in multiple neurological and neurodegenerative disorders (4–7).

KARs are tetrameric assemblies of GluK1-5 core subunits with GluK2/GluK5 heteromers, the most abundant postsynaptic KAR subunit combination (8). KAR trafficking and functional surface expression are tightly regulated through a variety of protein interactions and post-translational modifications (2, 9–11) and, in particular, the GluK2 subunit is post-translationally modified by phosphorylation (12–14), ubiquitination (15, 16), SUMOylation (17) and palmitoylation (18, 19) in its intracellular C-terminus.

SUMO1 is a 97-residue, 11 kDa protein that is covalently conjugated to specific lysine residues on target proteins to modify substrate function (4). GluK2 was the first of many synaptic membrane proteins shown to SUMOylated (17) and SUMOylation is now known to be involved in multiple neuronal signalling cascades and is strongly implicated in many neurological and neurodegenerative diseases. SUMOylation of GluK2 at K886 is required for agonist-induced internalisation of surface expressed GluK2-containing KARs and acts to regulate the complement of synaptic KARs (20). Importantly, PKC phosphorylation of S868 enhances GluK2 SUMOylation and is required for KAR LTD at MF-CA3 synapses (13, 14, 21).

In addition to mediating GluK2 SUMOylation, PKC phosphorylation has been shown to play a variety of roles in GluK2 surface delivery and activity-dependent trafficking, depending on the site and cellular location. For example, ER exit and secretory pathway trafficking of GluK2-containing KARs is restricted by phosphorylation of GluK2 at residues S846 and S868 by PKC (12, 22). Furthermore, PKC phosphorylation of S846 promotes endocytosis of GluK2-containing KARs, potentially by modulation of the interaction between GluK2 and 4.1 proteins (19), and PKC phosphorylation of S868 also participates in recycling of endocytosed GluK2 back to the plasma membrane (14). Thus, understanding the context-specific effects of GluK2 phosphorylation, and its interplay with other PTMs, represents an important objective in the field.

The dynamic and reversible addition of palmitate to proteins (palmitoylation) regulates the trafficking and surface stability of a wide variety of integral membrane proteins (23, 24). However, historically, palmitoylation has been difficult to study, so understanding of its targets and functions in neurons remains comparatively limited (25). Interestingly, however, in cell lines, all KAR subunits, except GluK1, can be palmitoylated at two conserved distal cytosolic cysteine residues (C858 and C871 in GluK2) (18, 19). Like phosphorylation and SUMOylation, palmitoylation is a highly dynamic, reversible post-translational process that can enhance the hydrophobicity of proteins to control target protein membrane association and clustering, especially at lipid rafts, in addition to regulating trafficking and protein-protein interactions (26, 27). Importantly, however, palmitoylation of GluK2 has not been demonstrated in neurons, and how interplay between palmitoylation, phosphorylation and SUMOylation coordinates KAR surface expression and activity-dependent endocytosis has not been investigated.

Here, we optimised an APEGS palmitoylation assay to investigate the interplay between palmitoylation, phosphorylation and SUMOylation of GluK2. We demonstrate that GluK2 is endogenously palmitoylated in neurons, and that GluK2 palmitoylation is reduced by agonist stimulation. Furthermore, using a non-palmitoylatable GluK2 mutant, we demonstrate that preventing GluK2 palmitoylation enhances PKC-mediated phosphorylation at S868, and SUMOylation at K886, and results in reduced KAR surface expression. Moreover, non-palmitoylatable GluK2 was insensitive to kainate-induced internalisation. Thus, our data support a model whereby agonist-induced depalmitoylation leads to PKC-mediated phosphorylation and subsequent SUMOylation of GluK2, to promote KAR endocytosis.

## Results

### GluK2 can be palmitoylated at C858 and C871 in HEK293T cells

The topology of GluK2, the amino acid sequence of the intracellular C-terminal domain and sites of key post-translational modifications are illustrated in Figure 1A. To measure GluK2 palmitoylation we modified the Acyl-PEGyl Exchange Gel Shift (APEGS) assay (28, 29). In this assay, cells are lysed in the presence of a reducing agent to break disulphide bonds, followed by treatment with the alkylating agent *N-*ethylmaleimide to ‘block’ thiol groups on free cysteines. Palmitoyl moieties are then cleaved from modified cysteines using hydroxylamine, and lysates treated with a high-molecular weight thiol-reactive mPEG-MAL-10K to specifically label formerly palmitoylated cysteines. Proteins modified by palmitoylation therefore exhibit a mass shift of ∼10kDa when analysed by Western blotting.

**Figure 1.**
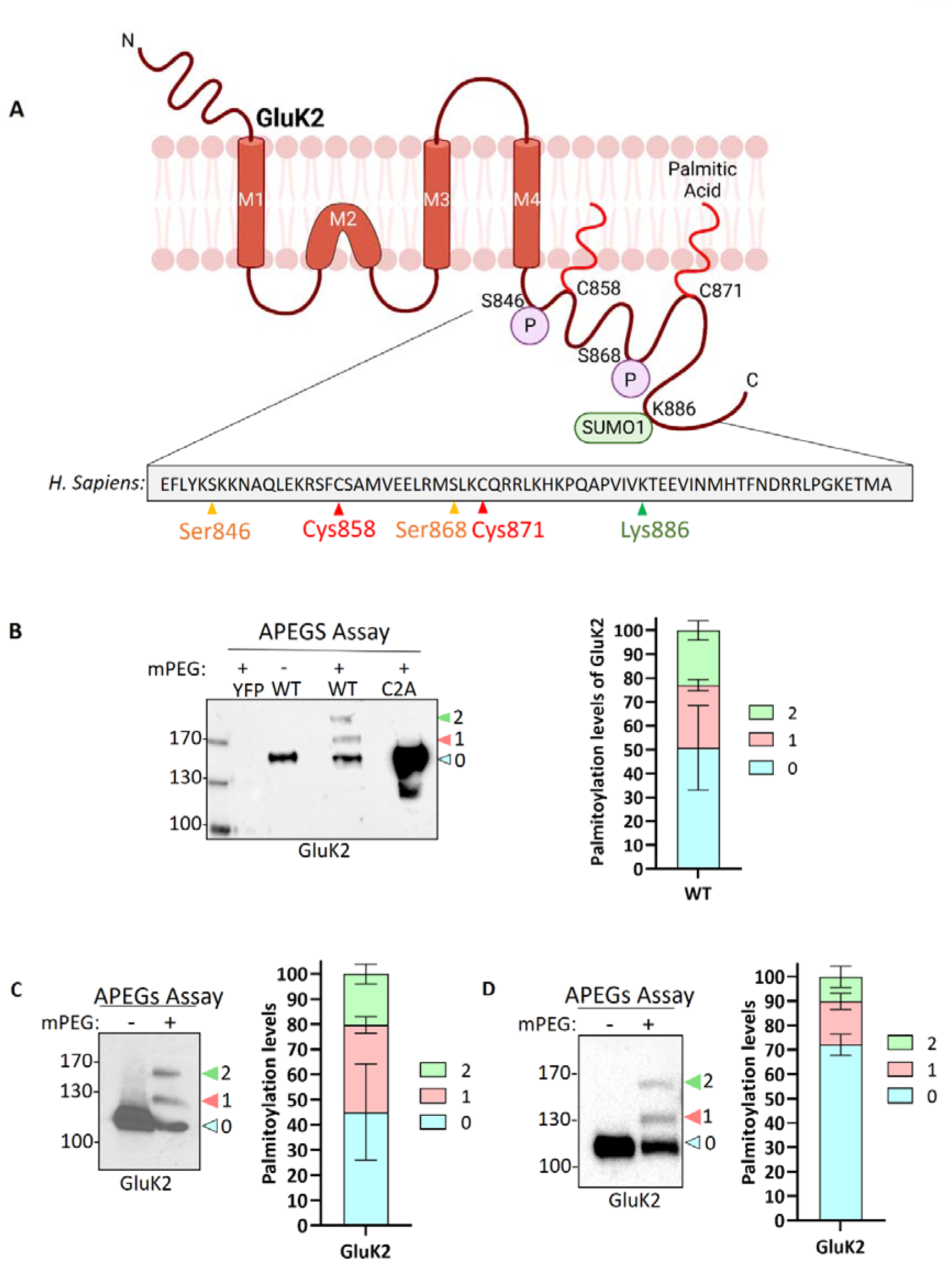
GluK2 is palmitoylated in HEK293T cells, mouse brain and cultured rat cortical neurons. A. The GluK2 subunit of KAR undergoes a complex array of post-translational modifications in its C-terminus, which include palmitoylation at cysteine residues 858 and 871, phosphorylation at serine residues 846 and 868, and SUMOylation at lysine residue 886. B. HEK293T cells were transfected with YFP, YFP-myc-GluK2 (WT), or the non-palmitoylated form of YFP-myc-GluK2 (C2A). An APEGs assay was then performed to evaluate the palmitoylation levels of recombinant homomeric GluK2. The results of this assay are reflected by a band shift, as a result of the addition of 10kDa mPEG-MAL molecules to the exposed cysteine residues. Data from the blot analysis indicates that GluK2 is palmitoylated at two distinct cysteine residues (C858/C871), since the double cysteine mutant did not exhibit any palmitoylation. In recombinant systems, approximately 50% of the total GluK2 protein exhibits palmitoylation at one or both of its cysteine residues. 27% of total GluK2 exhibits palmitoylation at a single cysteine residue, while an additional 23% of GluK2 was palmitoylated at both cysteine residues. In the analysis, “0” indicates non-palmitoylated GluK2, while “1” represents GluK2 that is palmitoylated at a single cysteine residue and “2” signifies GluK2 that is palmitoylated at both cysteine residues. N=3 separate experiments. Error bars = SEM. C. A mouse cortex was used as a source of lysate for the APEGs assay. The negative control lane (-mPEG) showed no evidence of palmitoylation, while endogenous GluK2 was observed to exist in two palmitoylated forms. The graph depicts the palmitoylation states of native GluK2, with 55% of the total population exhibiting palmitoylation. Approximately 20% of the total GluK2 is palmitoylated at a single cysteine residue, while 35% is palmitoylated at both cysteine residues. In the analysis, “0” indicates non-palmitoylated GluK2, while “1” represents GluK2 that is palmitoylated at a single cysteine residue and “2” signifies GluK2 that is palmitoylated at two cysteine residues. N=3 separate experiments. Error bars = SEM D. Rat cortical cells were plated and used for APEGs assay at DIV 19-21. 31% of total GluK2 was palmitoylated at either single (14.5%) or double (16.5%) cysteines. In the analysis, “0” indicates non-palmitoylated GluK2, while “1” represents GluK2 that is palmitoylated at a single cysteine residue and “2” signifies GluK2 that is palmitoylated at two cysteine residues. N=3 separate dissections. Error bars = SEM.

We first validated the APEGS assay by expressing YFP-myc-GluK2 wild-type (GluK2-WT) or a double cysteine mutant C858/C871A (GluK2-C2A), which has previously been reported to be non-palmitoylatable in cell lines (18, 19), in HEK293T cells. As shown in Figure 1B, GluK2-WT showed two higher molecular weight bands corresponding to singly and doubly palmitoylated GluK2. These bands were absent in the negative control lane in which mPEG-MAL-10K was omitted, confirming them as genuine mPEG adducts. Consistent with previous reports (18, 19), no higher molecular weight bands were present in the GluK2-C2A mutant, even though the expression of the latter construct was much greater, confirming C858 and C871 as the only palmitoylated cysteines in GluK2. Quantification of our data shows that ∼50% of the GluK2 expressed in HEK293T cells is not palmitoylated, whereas ∼27% is singly palmitoylated and ∼23% is doubly palmitoylated (Figure 1C).

### Endogenous GluK2 is palmitoylated in neurons

Having established that recombinant GluK2 is palmitoylated in HEK293T cells we next used the APEGS assay to investigate palmitoylation of endogenous GluK2 in mouse cortex (Figure 1C) and cultured rat cortical neurons (Figure 1D). Consistent with the HEK293T cell data, in both systems we detected two palmitoylated forms of endogenous GluK2. In mouse cortex, ∼35% of endogenous GluK2 is singly and ∼20% doubly palmitoylated, while in DIV 19-21 rat cortical neurons ∼14.5% was singly and ∼16.5% doubly palmitoylated. Together, these data confirm palmitoylation of endogenous GluK2.

### Kainate stimulation decreases GluK2 palmitoylation

Having confirmed palmitoylation of endogenous GluK2, we next wondered whether palmitoylation was dynamically regulated by KAR activation. We therefore pre-incubated neurons with 2µM TTX and 40µM GYKI53655, to block spontaneous activity and prevent KA-induced activation of AMPARs, respectively, and then stimulated neurons with 20µM KA for 5 mins. Interestingly, KAR activation led to a decrease in GluK2 double palmitoylation (Figure 2 A, B), suggesting that agonist-stimulation leads to a rapid depalmitoylation of GluK2.

**Figure 2.**
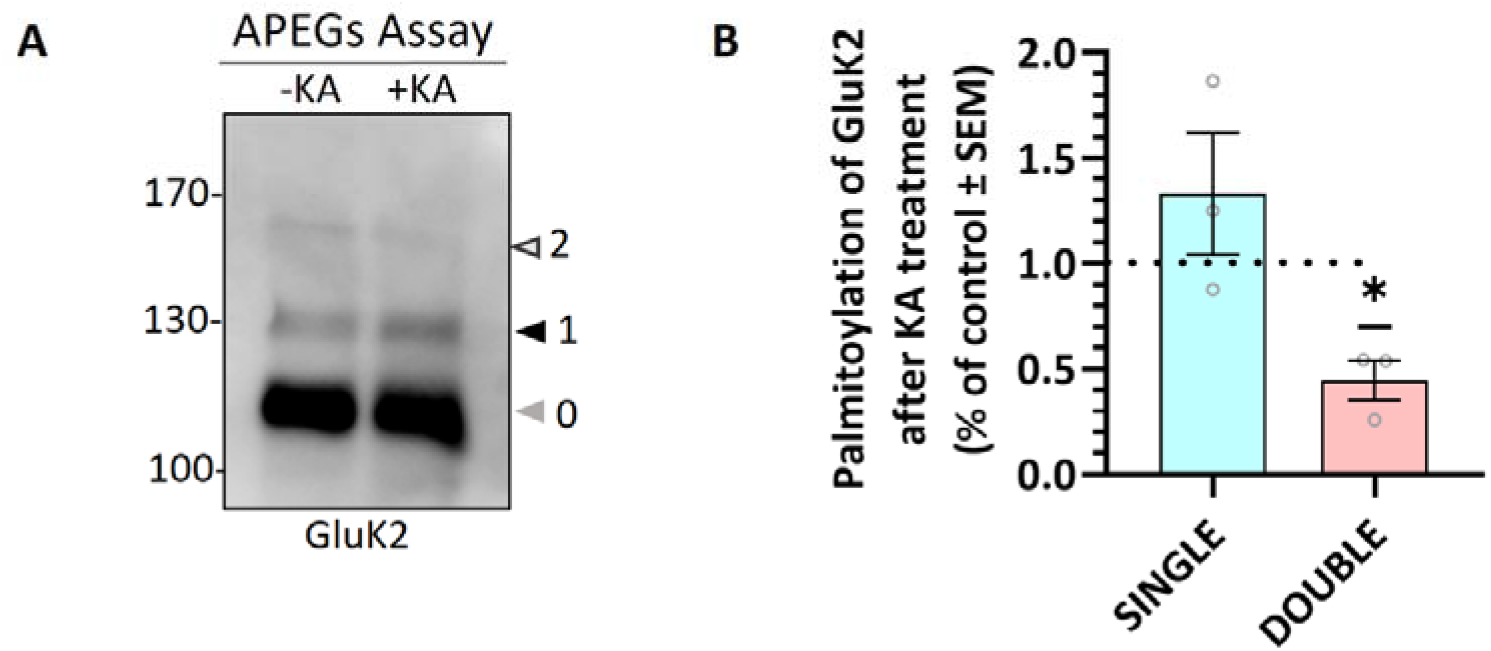
GluK2 palmitoylation decreases with KA stimulation. DIV 19-21 cultured cortical neurons were stimulated with 20µM KA for 5 mins and lysates were subjected to the APEGS assay. The graph shows the comparison of single and double palmitoylation in control and KA-induced cells. Control single and double palmitoylation values were set as one and KA-induced values are shown as a percentage of the control. There was a significant decrease in the level of double palmitoylated GluK2 in KA-induced neurons compared to the control. N=3 independent dissections. One sample t-test (\**p*<0.05, \*\**p*<0.001, \*\*\**p*<0.0001). Error bars = SEM.

### GluK2 is palmitoylated at C858 and C871 in cultured neurons

To further investigate GluK2 palmitoylation and its interplay with other post-translational modifications, we next used a lentiviral GluK2 shRNA knockdown (KD) and replacement strategy in rat cortical neuronal cultures (30). Lentiviral constructs knocked down endogenous GluK2 and simultaneously expressed shRNA-insensitive YFP-myc-tagged GluK2 variants. Specifically, we replaced endogenous GluK2 with either GluK2-WT, non-palmitoylatable GluK2-C2A or a phosphorylation-deficient GluK2-S2A (S846/868A), which lacks two previously characterised PKC phosphorylation sites in the GluK2 C-terminus (12). Neurons were infected with KD-replacement lentivirus at DIV 14 and after a further 7 days cells were lysed and Western blotted to ensure that each YFP-myc-GluK2 mutant was expressed at similar levels and the expected molecular weight (∼150 kDa) (Figure 3A).

**Figure 3.**
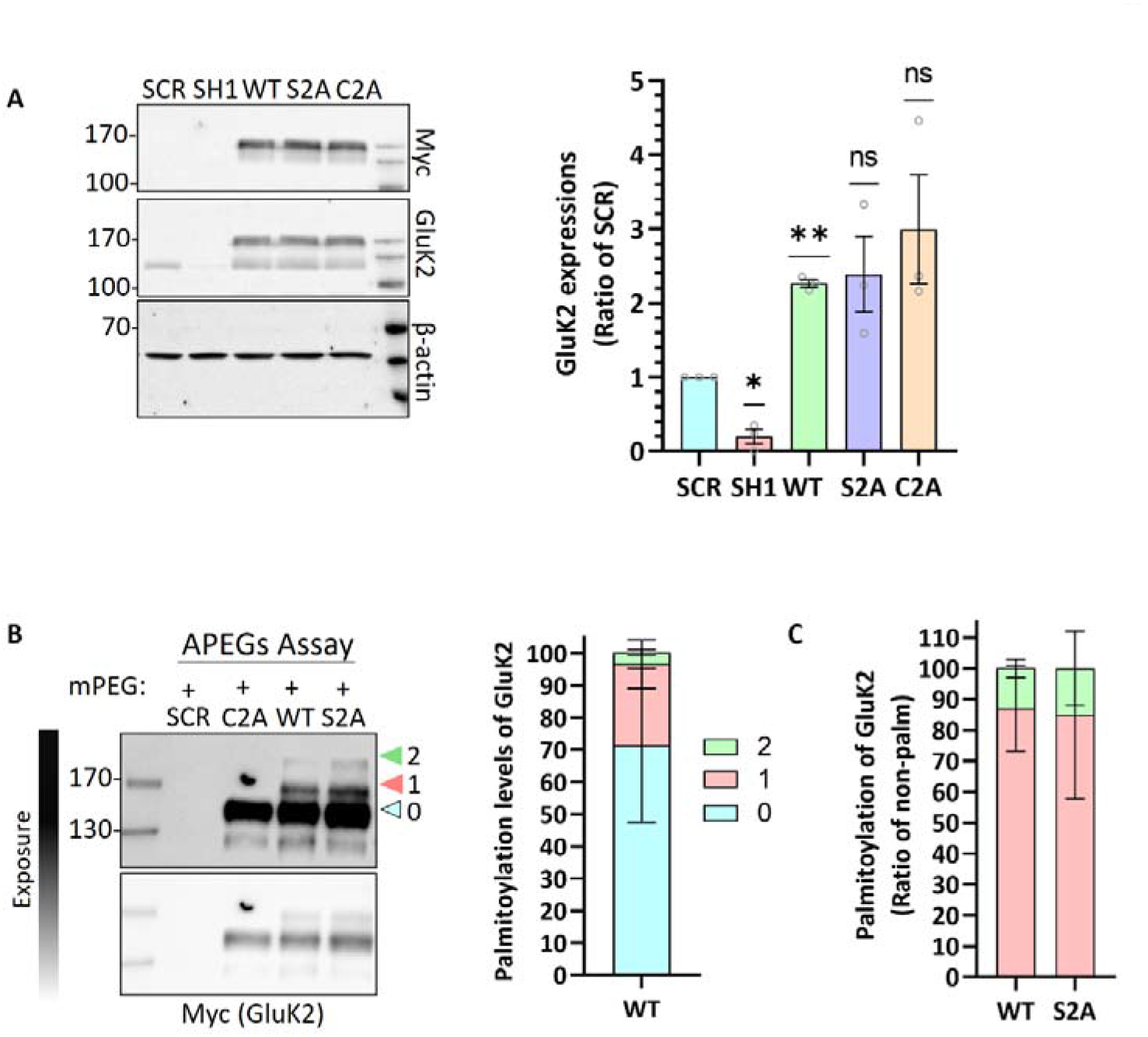
GluK2 is palmitoylated at C858 and C871 in cultured cortical neurons. A. DIV 14 rat cortical neurons were infected with GluK2-expressing lentiviruses: control (GFP), wild-type (YFP-myc-GluK2), non-phosphorylatable (YFP-myc-GluK2-S2A) or non-palmitoylatable (YFP-myc-GluK2-C2A). The graph shows expression levels of the different GluK2 mutants. Data was normalised to beta-actin. One sample t-test (\**p<*0.05, \*\**p*<0.01). N=3 dissections. Error bars = SEM. B. DIV 14 rat cortical neurons were infected with GluK2-expressing lentiviruses: control (GFP), wild-type (YFP-myc-GluK2), non-phosphorylatable (YFP-myc-GluK2-S2A) or non-palmitoylatable (YFP-myc-GluK2-C2A). The palmitoylation levels of the recombinant GluK2 mutants were analysed using the APEGS assay. The results confirm that the double cysteine mutant, C858A/C871A is not palmitoylated in neurons. The graph depicts the palmitoylation states of native GluK2, with 30% of the total population exhibiting palmitoylation. Approximately 26% of the total GluK2 population is palmitoylated at a single cysteine residue, while 4% of the population is palmitoylated at both cysteine residues. In the analysis, “0” indicates non-palmitoylated GluK2, while “1” represents GluK2 that is palmitoylated at a single cysteine residue and “2” signifies GluK2 that is palmitoylated at two cysteine residues. N=3 separate dissections. Error

We then assessed the palmitoylation of GluK2-WT, GluK2-C2A and GluK2-S2A using the APEGs assay. As expected, GluK2-WT showed bands corresponding to single and double palmitoylation, but these bands were absent for GluK2-C2A, confirming in neurons that only C858 and C871A can be palmitoylated. Consistent with the overall levels of palmitoylation observed for endogenous GluK2 in cultured neurons, ∼30% of total YFP-myc-tagged GluK2-WT was palmitoylated (Figure 3B). Palmitoylation levels of the PKC phospho-null mutant GluK2-S2A were similar to GluK2-WT, suggesting that phosphorylation of GluK2 does not play a major role in regulating GluK2 palmitoylation (Figure 3C).

### Non-palmitoylatable GluK2 shows enhanced PKC phosphorylation at S868

We have shown previously that agonist stimulation of GluK2 promotes PKC-mediated phosphorylation at S868, leading to enhanced SUMOylation at K886 and receptor endocytosis (13, 14, 17). Interestingly, it has been reported previously using recombinant proteins that non-palmitoylatable GluK2 is a better substrate for phosphorylation by PKC than WT GluK2 (18). Since KA stimulation leads to a reduction in GluK2 palmitoylation, we therefore wondered whether receptor depalmitoylation promotes PKC phosphorylation and subsequent endocytosis. To examine this directly, we used phos-tag gel electrophoresis (31, 32) to analyse phosphorylation of non-palmitoylatable GluK2.

Endogenous GluK2 was knocked down in cortical neurons and replaced with YFP-myc-tagged GluK2-WT, non-palmitoylatable GluK2-C2A or, as a negative control for PKC phosphorylation, GluK2-S2A (S846/868A). Neurons were infected with the relevant lentivirus at DIV 14 and lysed at DIV 21 and samples were subjected to phos-tag gel electrophoresis and Western blotting. Under basal conditions no band shifts indicating phosphorylated GluK2 were detected in GluK2-WT or PKC phospho-null GluK2-S2A. Interestingly, however, we observed clear phosphorylation of the non-palmitoylatable GluK2-C2A mutant (Figure 4A), suggesting this mutant may be more phosphorylated than its WT counterpart.

**Figure 4.**
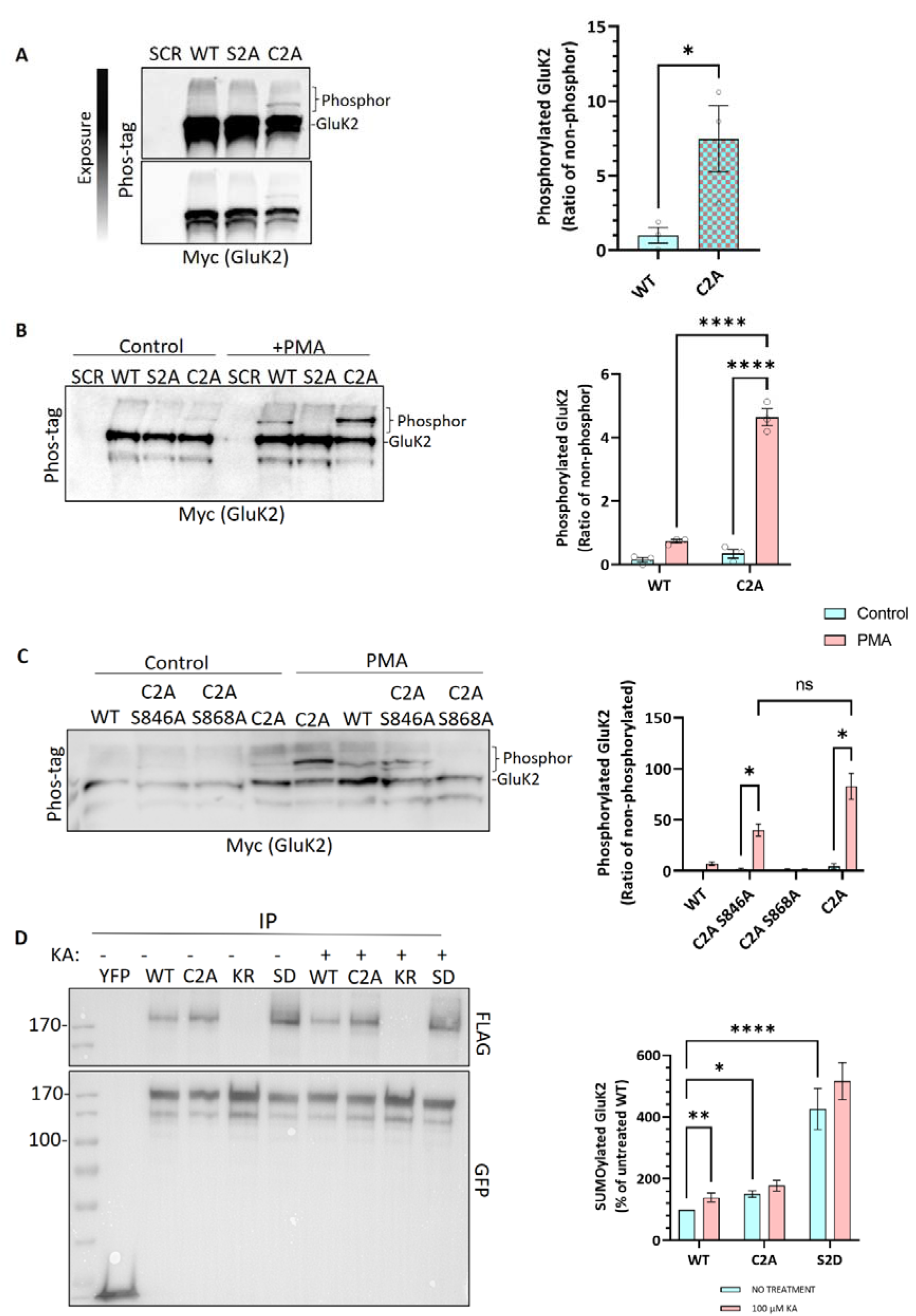
Non-palmitoylated GluK2 is a better substrate for protein kinase C than wild-type GluK2. A. Cultured rat cortical neurons were infected with the indicated lentiviruses: GFP, YFP-myc-GluK2 (WT), non-phosphorylatable YFP-myc-GluK2 (S2A) or non-palmitoylatable YFP-myc-GluK2 (C2A) at DIV 14. After 7 days of incubation, cells were lysed with 2x sample buffer and run on a phos-tag gel followed by Western blotting for myc. Band shifts indicate GluK2 phosphorylation for the C2A mutant, but not WT or S2A. Phosphorylated GluK2 was first normalised to non-phosphorylated GluK2 and C2A expressed as a proportion of the WT. Student t-test (\**p<* 0.05, \*\**p*<0.01, \*\*\**p*<0.001). B. As for A except that after 7 days of incubation cells were incubated with 1µM PMA for 20 min and lysed with 2x sample buffer, sonicated, and run on a phos-tag gel. Western blots were then probed for myc. Phosphorylated GluK2 bands in WT and C2A were first normalised to unconjugated GluK2. Two-way ANOVA with Tukey’s multiple comparisons test (\**p<*0.05, \*\**p*<0.01, \*\*\**p*<0.001, \*\*\*\**p*<0.0001). N=3 independent dissections. Error bars = SEM. C. Cultured rat cortical neurons were infected with the indicated lentiviruses: GFP, YFP-myc-GluK2 (WT), single PKC phosphorylation-deficient and non-palmitoylated YFP-myc-GluK2 (C2A-S846A or C2A-S868A), and non-palmitoylated YFP-myc-GluK2 (C2A) at DIV 14. After 7 days of incubation, cells were incubated with 1µM PMA for 20 min and lysed with 2x sample buffer, sonicated, and run on a phos-tag gel. Western blot was probed for myc. Two-way ANOVA with Tukey’s multiple comparisons test (\**p<* 0.05, \*\**p*<0.01, \*\*\**p*<0.001, \*\*\*\**p*<0.0001). N=3 independent dissections. Error bars = SEM. D. Preventing palmitoylation of GluK2 mimics agonist-induced SUMOylation. HEK293T cells were transfected with either YFP or the indicated YFP-myc-GluK2 constructs, along with FLAG-tagged SUMO1 and FLAG-tagged Ubc9. 48 hours post-transfection, cells were stimulated with 100µM kainate for 20 minutes. GluK2 was immunoprecipitated (IP) using GFP-trap and the levels of SUMOylated GluK2 were detected in the different conditions by Western blotting using an anti-FLAG antibody. SUMOylated GluK2 signal was first normalised to the amount of YFP-GluK2 pulled down before being normalised to the untreated WT. To compare the different mutants (untreated and treated) with the WT (untreated and treated), Two-way ANOVA with Tukey’s multiple comparisons test was applied (\**p<*0.05, \*\**p*<0.01, \*\*\**p*<0.001, \*\*\*\**p*<0.0001). N=6 separate experiments. Error Bars = SEM.

To directly investigate the effect of GluK2 palmitoylation on PKC phosphorylation neurons were incubated with the PKC activator phorbol 12-myristate 13-acetate (PMA; 1µM for 20 mins) before lysis and subjected to phos-tag gel electrophoresis followed by Western blotting (Figure 4B). As expected, no PKC phosphorylation was detected for GluK2-S2A, but both GluK2-WT and GluK2-C2A showed enhanced phosphorylation in response to PKC activation. Furthermore, the non-palmitoylatable GluK2-C2A was phosphorylated to a ∼5-fold greater extent than GluK2-WT, confirming that non-palmitoylatable GluK2 is a better substrate for PKC phosphorylation in neurons, and suggesting that agonist-induced depalmitoylation of GluK2 may lead to phosphorylation by PKC.

We next sought to examine which PKC site on GluK2 shows enhanced phosphorylation in the absence of receptor palmitoylation. We therefore generated non-palmitoylatable mutants of GluK2 in which single PKC phosphorylation sites (S846 or S868) were mutated to alanine. Cortical neurons were infected with knockdown-rescue lentiviruses expressing YFP-myc-tagged GluK2-WT, GluK2-C2A, or the non-palmitoylatable single PKC phosphorylation mutants GluK2-C2A/S846A or GluK2-C2A/S868A at DIV 14. After 7 days, cells were incubated with 1µM PMA for 20 min, subjected to phos-tag gel electrophoresis and blots were probed for YFP-myc-GluK2 (Figure 4C). Interestingly, while GluK2-WT, GluK2-C2A and GluK2-C2A/S846A underwent phosphorylation in response to PKC activation, GluK2-C2A/S868A did not, suggesting that the enhanced phosphorylation of the non-palmitoylatable GluK2 mutant occurs S868. Together, these data suggest that in the absence of palmitoylation GluK2 is more readily phosphorylated by PKC at S868.

### Non-palmitoylatable GluK2 exhibits enhanced SUMOylation

Since the non-palmitoylatable mutant of GluK2 exhibits enhanced PKC phosphorylation at S868, and phosphorylation at this site has been demonstrated to enhance GluK2 SUMOylation (13, 14), we next examined SUMOylation of the non-palmitoylatable GluK2-C2A. We transfected HEK293T cells with YFP-myc-tagged GluK2-WT, non-palmitoylatable GluK2-C2A or, as a negative control, non-SUMOylatable GluK2-K886R. To aid detection of SUMOylation, cells were also co-transfected with FLAG-SUMO1 and FLAG-Ubc9. 48 hours post-transfection, cells were stimulated with 100µM KA for 20 minutes before lysis and GluK2 SUMOylation was quantified by GFP-trap pulldown, to isolate YFP-myc-tagged GluK2, followed by blotting for FLAG-SUMO1 (Figure 4D). Interestingly, under basal conditions there was greater SUMOylation of non-palmitoylatable GluK2-C2A than GluK2-WT, consistent with enhanced phosphorylation of this mutant at S868. Furthermore, while kainate stimulation evoked a significant increase in SUMOylation of GluK2-WT, SUMOylation of GluK2-C2A was not further enhanced. Taken together, these data suggest that enhanced PKC phosphorylation of non-palmitoylated GluK2 at S868 enhances GluK2 SUMOylation, and support a model whereby agonist-induced depalmitoylation of GluK2 is required to illicit receptor SUMOylation in response to KA.

### Decreased surface expression of non-palmitoylatable GluK2

We next infected neurons with knockdown-rescue lentiviruses expressing YFP-myc-tagged GluK2-WT or GluK2-C2A at DIV 14-15. 7 days later, neurons were treated with 2µM tetrodotoxin (TTX) and 40µM GYKI53655 for 30 min, then stimulated with 10µM KA for 20 min. Surface expressed proteins were then labelled with membrane impermeant biotin, followed by isolation of biotinylated proteins with streptavidin beads and Western blotting (Figure 5A). Consistent with the SUMOylation data, surface expression of GluK2-C2A was significantly reduced compared to GluK2-WT and was not further reduced upon KA stimulation (Figure 5B).

**Figure 5.**
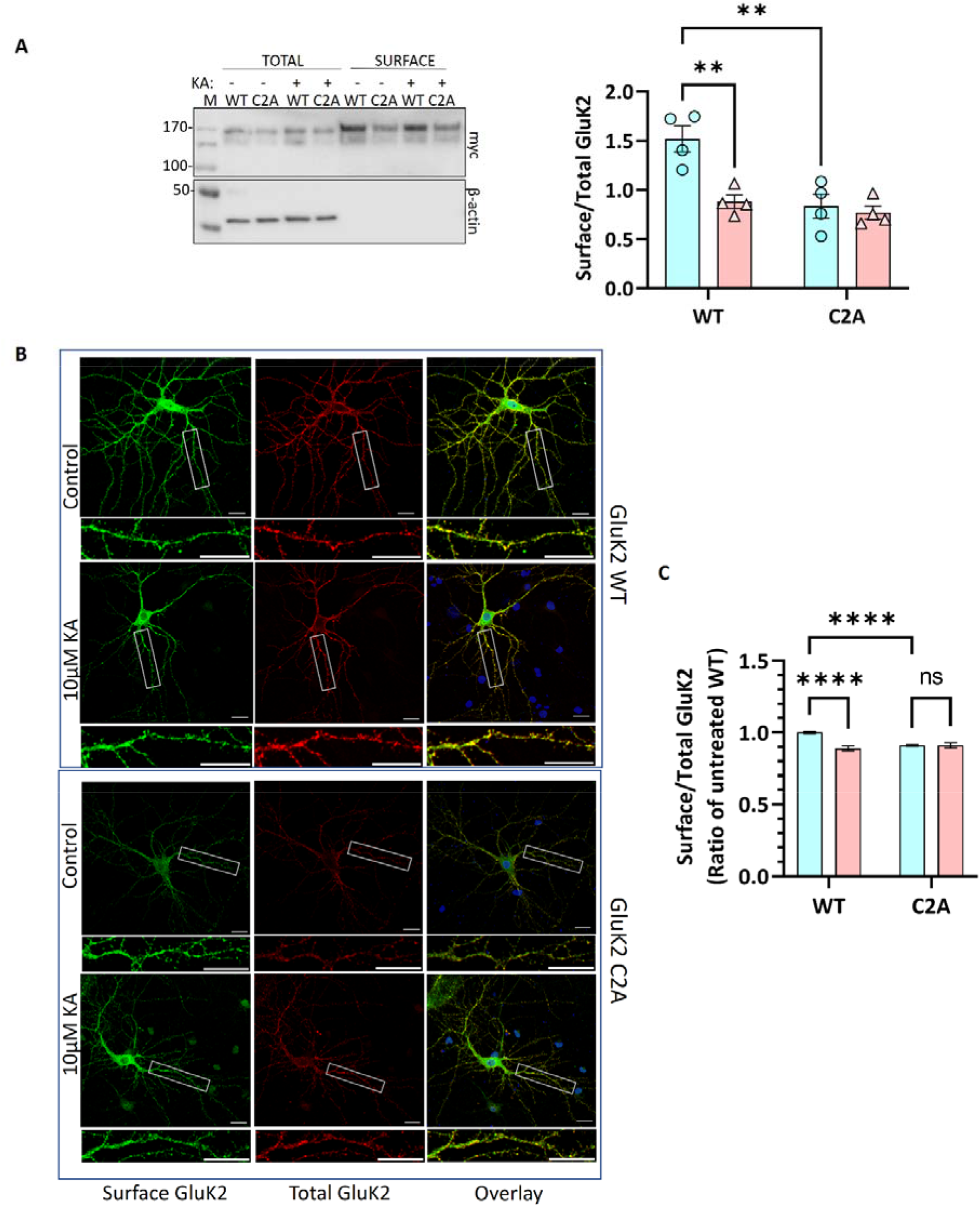
A. Cultured rat cortical neurons were infected with lentiviruses as indicated: SCR, YFP-myc-GluK2 (WT) or double non-palmitoylated YFP-myc-GluK2 (C2A) at DIV 14. After 7 days of incubation neurons were pre-treated with 2µM TTX and 40µM GYKI53655 for 30 min before being treated with 10µM kainate for 20 min. Surface biotinylation experiments were performed followed by streptavidin pulldown and Western blotting. Surface levels of GluK2 were normalised to total levels. Two-way ANOVA with Tukey’s multiple comparisons test (\**p*< 0.05, \*\**p*<0.01, \*\*\**p*<0.001, \*\*\*\**p*<0.0001). N=4 dissections. Error Bars = SEM. B. Preventing palmitoylation of GluK2 reduces surface expression and occludes agonist-induced internalisation. Representative images of hippocampal neurons transfected with YFP-myc-tagged GluK2 (WT or non-palmitoylatable C2A) at DIV 9. At DIV 14 neurons were pre-treated with 2µM TTX and 40µM GYKI53655 for 30 min before being treated with 10µM kainate for 20 min. Live labelling (using anti-GFP antibody followed by Alexa 647) was used to label the surface expressed GluK2 receptors (red). The total receptors were immunostained by anti-GFP followed by Cy2 (green) after the red staining of the surface receptors as described in the methods. 5-15 cells per condition per experiment (a total of 3 independent experiments) were analysed using ImageJ Fiji software. C. Quantification of the fluorescence imaging data shown in B). The results are presented as the surface/total ratio of GluK2 expressed as a ratio of the untreated WT in both the dendrites and soma. Data presented as a percentage of the untreated WT. Two-way ANOVA with Tukey’s multiple comparisons test (\**p<*0.05, \*\**p*<0.01, \*\*\**p*<0.001, \*\*\*\**p*<0.0001). Scale bars: 20 µm (main panel) and 5 µm (magnification panel). 5-15 cells/condition/experiment. N=3 independent experiments. Error Bar = SEM.

**Figure 6.**
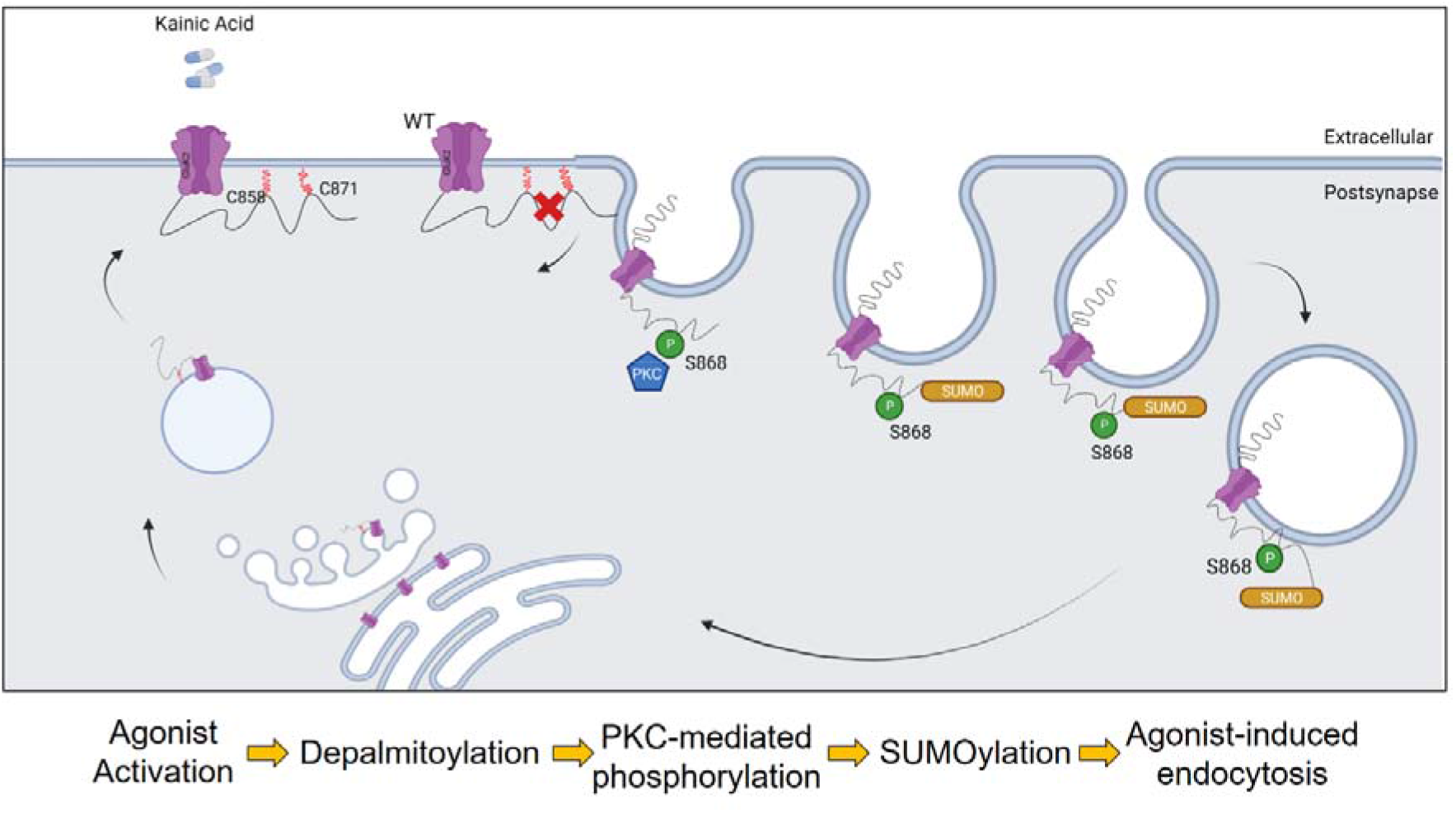
Schematic model. The C-terminus of the KAR subunit GluK2 has two cysteine residues (C858 and C871) that are substrates for palmitoylation. Depalmitoylation of GluK2 occurs in response to agonist activation, and leads to phosphorylation of the nearby serine, S868, by PKC. PKC phosphorylation then promotes SUMO1 conjugation to GluK2 at K886, resulting in receptor internalisation.

To further confirm these findings, we used confocal microscopy to compare the surface expression and agonist-induced endocytosis of YFP-myc-tagged GluK2-WT and non-palmitoylatable GluK2-C2A in cultured rat hippocampal neurons. Neurons were transfected at DIV 9-10 and at DIV 14-15 treated with 2µM tetrodotoxin (TTX) and 40µM GYKI53655 for 30 min to prevent spontaneous synaptic activity, before stimulation with 10µM kainate for 20 min to induce receptor internalisation (13, 17). Neurons were then live labelled with anti-GFP antibodies to measure YFP-myc-GluK2-WT and -C2A surface expression, followed by fixation, permeabilisation, and staining to assess total YFP-myc-GluK2 (Figure 5D). Interestingly, and consistent with the surface biotinylation data, the surface expression of GluK2-C2A was significantly reduced compared to GluK2-WT under basal conditions. Moreover, while surface expression of GluK2-WT was significantly decreased upon KA stimulation (Figure 5C), GluK2-C2A was not, suggesting that preventing GluK2 palmitoylation occludes KA-induced receptor endocytosis.

Taken together, our data support a model whereby activity-dependent depalmitoylation of GluK2 leads to enhanced PKC-mediated phosphorylation at S868, receptor SUMOylation, and KAR endocytosis.

## Discussion

Here we adapted and optimised an APEGS assay to investigate GluK2 kainate receptor subunit palmitoylation and its interplay with other PTMs. The APEGS assay allows us to explore the palmitoylation pattern of low abundance palmitoylated proteins under native conditions with high sensitivity. Previous radiolabelled [^3^H]-palmitate assays in heterologous systems indicated that GluK2 can be palmitoylated at C-terminal cysteine residues C858 and C871. In those studies, both C858A and C871A mutants showed reduced palmitoylation, but it was unclear if both cysteines were simultaneously palmitoylated (18, 19). We engineered a construct in which both C858 and C871 were mutated to alanine, GluK2-C2A. For the first time we show that while GluK2-C2A exhibits no palmitoylation, WT GluK2 is palmitoylated at both cysteine residues simultaneously when recombinantly expressed in HEK293T cells, and endogenously in cortical neurons and brain. More importantly, we show that kainate stimulation causes depalmitoylation of endogenous GluK2, suggesting that dynamic depalmitoylation may contribute to activity-dependent KAR trafficking.

We then used a lentiviral approach to knock down endogenous GluK2 in cortical neurons and replace it with YFP-myc-GluK2-WT or YFP-myc-GluK2-C2A and observed, using phos-tag electrophoresis, that preventing palmitoylation of GluK2 enhanced PKC phosphorylation. An increase in GluK2 phosphorylation of non-palmitoylated GluK2 in heterologous systems has also been reported previously (18), however had not been demonstrated in neurons. Here, we extended these findings by showing that preventing palmitoylation of GluK2 specifically enhances phosphorylation of the S868 PKC phosphorylation side, further supporting the proposal that interplay between palmitoylation and phosphorylation act to fine-tune KAR trafficking. Potentially, depalmitoylation acts to ‘release’ the C-terminal domain of GluK2 from the inner membrane leaflet to allow access for PKC phosphorylation.

Since kainate stimulation leads to depalmitoylation of GluK2 and PKC phosphorylation at S868, we then examined SUMOylation of non-palmitoylatable GluK2. Consistent with previous work showing that S868 phosphorylation promotes GluK2 SUMOylation (13, 14), non-palmitoylatable GluK2 exhibited enhanced SUMOylation, which was not further increased by agonist treatment, suggesting GluK2 depalmitoylation represents a permissive step in mediating receptor phosphorylation, SUMOylation and subsequent endocytosis. In agreement with this model, GluK2-C2A exhibited lower surface expression levels compared to the GluK2-WT under basal conditions and was insensitive to agonist-induced endocytosis. Thus, our data are consistent with activity-dependent interplay between palmitoylation and SUMOylation representing a requirement for agonist-induced GluK2 endocytosis.

The regulation of protein-protein interactions and synaptic vesicle dynamics by the coordinated, often bidirectional, actions of palmitoylation and phosphorylation has been reported to play a role in striatal dopamine release (33), neuronal outgrowth (34) and synapsin 1-mediated clustering of synaptic vesicles (35). Directly relevant to our work, phosphorylation of GluK2 has been reported to eliminate the interaction of GluK2 with 4.1N protein, thereby decreasing KAR surface expression. Conversely, the same study reported that GluK2 palmitoylation promotes interaction with 4.1N, stabilising KARs at the cell surface (36). Taken together, our results suggest that activity-dependent depalmitoylation, followed by consequent S868 phosphorylation and SUMOylation, may act collectively to mediate uncoupling of GluK2 from 4.1N, to facilitate receptor endocytosis in response to agonist activation.

Here we show that depalmitoylation of GluK2 leads to SUMO1-dependent KAR endocytosis. Thus, we propose that agonist-induced PKC-mediated phosphorylation of GluK2 acts as an intermediate that ‘links’ depalmitoylation to SUMOylation, and consequent endocytosis. Together, these findings provide new insights into the molecular mechanisms that regulate KAR trafficking and expression in response to neuronal activity.

## Materials and Methods

### Drugs

| Drugs | Concentration<br>(μM) | Diluted<br>in | Supplier | Catalogue Number |
| --- | --- | --- | --- | --- |
| **GYKI**<br>**53655** | 40 | ddH<sub>2</sub>O | Tocris | 2555 |
| **Kainate**<br>**(KA)** | 10 | ddH<sub>2</sub>O | Tocris | 0222 |
| **TTX** | 2 | ddH<sub>2</sub>O | Tocris | 1069 |

### Primary Antibodies

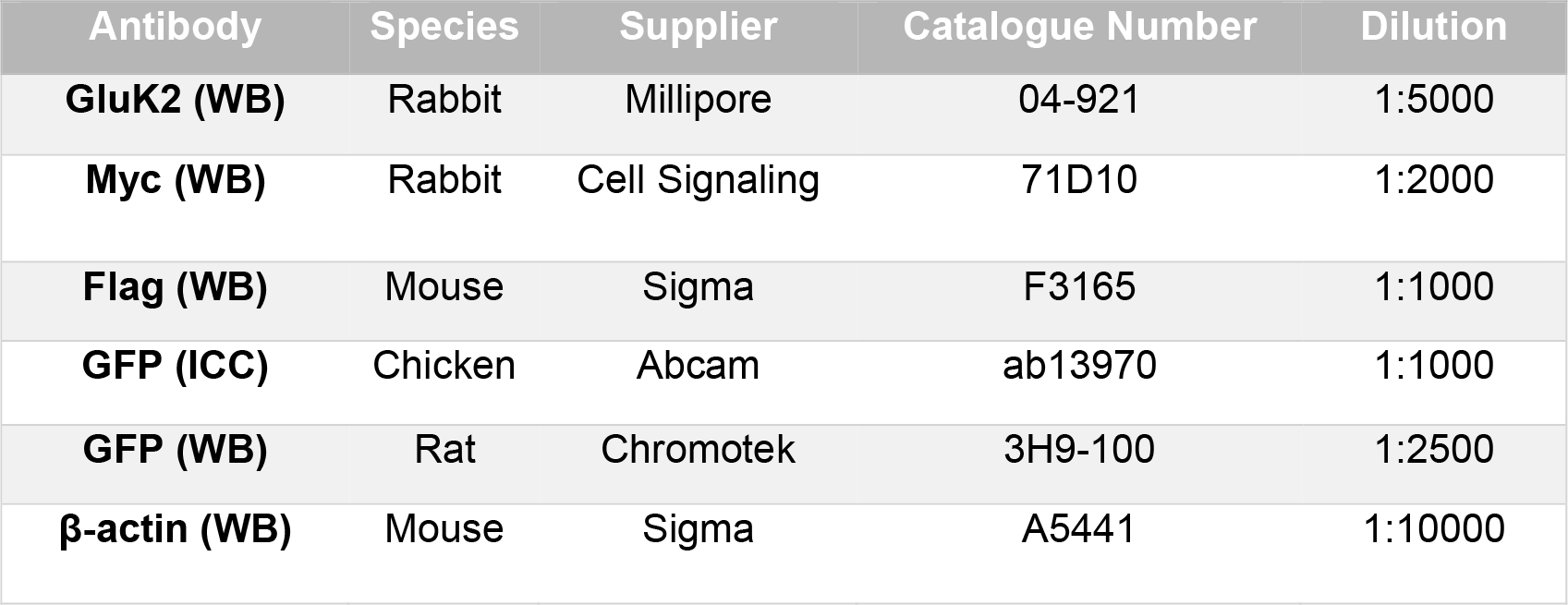

### Transfection of HEK293T Cells

Human Embryonic Kidney 293T (HEK293T) cells were cultured in high-glucose DMEM containing glutamine (Sigma) supplemented with 10% FBS and 2% Penicillin-Streptomycin. HEK293T cells were plated for transfection in 60mm dishes the day before transfection. YFP-myc-tagged GluK2 constructs, or GFP control, (2.5µg each DNA) were added to 500µl plain DMEM, followed by addition of Lipofectamine (1.5µl per µg DNA), vortexed briefly, and incubated for 20 min. The media on the cells was then aspirated and replaced with DMEM containing 10% FBS, and the DNA-Lipofectamine mixture added dropwise to the cells, and the transfected cells incubated for 48 hours before use.

### Neuronal Cell Cultures

Hippocampal cultures derived from E18 Han Wistar rats were prepared as described previously (13). The cortical cells were plated at 6-well dishes (500k per well) for biochemistry and hippocampal cells were plated on coverslips (100k per coverslip) for imaging. Cells were incubated for a period of 19-21 days before use. Initial the neuronal plating media consisted of Neurobasal (Gibco) supplemented with 10% Horse Serum, 2% B27, 2mM Glutamax, and Penicillin-Streptomycin. After 2 hours, the media was switched to feeding media lacking horse serum.

### Transduction with Lentivirus

To conduct GluK2 knockdown experiments, shRNA sequences specifically targeting GluK2 were inserted into a modified pXLG3-GFP vector (30), which was controlled by an H1 promoter. The shRNA target sequences used were as follows:

Control, non-targeting shRNA: AATTCTCCGAACGTGTCAC

GluK2-targeting shRNA: GCCGTTTATGACACTTGGA

The viral particles containing these shRNA constructs were generated in HEK293T cells as described previously (30). Subsequently, the harvested viral particles were added to cortical neurons at 12-14 days in vitro (DIV) and allowed to incubate for 7 days before being used for further experiments.

### Sustained KA stimulation

Hippocampal neurons were plated in a 6-well dish with a density of 500,000 cells per well. The cells were incubated at 37°C until reaching a developmental stage of DIV 19-21 DIV. On the experiment day, the cells were subjected to pre-treatment of 1µM TTX (Tocris) and 40µM GYKI 53655 (Tocris) to the culture medium for 30 min. Following pre-treatment, KA incubation (10µM for 20 min at 37°C) was performed in Earle’s Buffer (140mM NaCl, 5mM KCl, 1.8mM CaCl2, 0.8mM MgCl2, 25mM HEPES, and 5mM D-glucose pH of 7.4). Control cells were treated with a vehicle instead of KA.

### Neuronal Surface Biotinylation

After KA treatment, surface proteins on the cell were labelled using membrane impermeant Sulfo-NHS-SS-Biotin (ThermoFisher). The biotinylation process was carried out on ice, following a previously described protocol (37). All wash steps involved the use of ice-cold Earle’s Buffer to maintain the low temperature during the procedure. Subsequently, biotin-labelled surface proteins were isolated by streptavidin pulldown (37). Samples were the run on SDS-PAGE followed by Western blot with anti-myc and anti-β-actin antibodies.

### Detection of post-translational modifications

HEK293T cells or neurons were transfected or infected, respectively, with wild-type or point mutants of YFP-myc tagged GluK2 (non-palmitoylatable, C858A/C871A; non-SUMOylatable, K886R; phospho-null, S846A/S868A; phospho-mimetic, S846D/S868D; Non-palmitoylated, non-phosphorylated single mutants, C858A/C871A/S846A and C858A/C871A/S868A).

#### SUMOylation

HEK293T cells were transfected with GFP or YFP-myc-GluK2 constructs and Flag-SUMO1 and Flag-Ubc9. 48 hours after transfection, cells were stimulated with 100µM KA and lysed in lysis buffer (20mM Tris-HCl (pH 7.4), 137mM NaCl, 2mM sodium pyrophosphate, 2mM EDTA, 1% Triton-X 100, 0.1% SDS, 25mM β-glycerophosphate, 10% glycerol, 20mM N-ethylmaleimide (NEM, freshly prepared), and 1x complete protease inhibitors). Lysates were sonicated briefly and incubated on ice for 20 minutes, before being cleared by centrifugation at 16000g for 20 minutes at 4°C. The supernatant was then taken, and GluK2 was pulled down using GFP-Trap A beads (ChromoTek) overnight at 4°C. After three washes in lysis buffer, bound proteins were detected by SDS-PAGE followed by Western blotting using anti-Flag and anti-GFP antibodies.

#### Acyl-PEGyl Exchange Gel Shift (APEGS) Assay

We combined two previously published protocols to optimise the APEGS assay for GluK2 (28, 38, 39). Briefly, the APEGS assay comprises four steps: 1) disruption of disulphide bonds; 2) blocking free cysteine residues; 3) cleaving palmitoyl-thioester linkages with hydroxylamine; 4) labelling previously palmitoylated but now newly exposed cysteine residues with mPEG-MAL-10k. This results in a ∼10kDa increase in mass for each palmitoylated cysteine on SDS-PAGE (38).

Neurons were homogenised in PBS (Gibco) containing 4% SDS, 5mM EDTA, and protease inhibitors (cOmplete). After sonication, supernatant proteins (2 mg/ml) were reduced with 25mM TCEP (ThermoFisher) for 1 h at 55°C, followed by alkylation of free cysteine residues with 50mM NEM (Sigma) overnight at RT. The samples were then subjected to methanol/chloroform precipitation and the precipitate spun down, washed, and the protein pellet resuspended and incubated in 2M hydroxylamine (NH_2_OH) (Sigma), pH 7.0, 5mM EDTA, 0.2% (w/v) Triton X-100 for 1 h at 37°C to cleave palmitoylation thioester bonds. After a second methanol/chloroform precipitation samples were incubated with 7mM mPEG-

MAL10k (Sigma) for 2h at RT to label newly exposed cysteinyl thiols. As a negative control, mPEG was omitted. Following a third methanol/chloroform precipitation the protein pellet was resuspended with SDS-sample buffer (without β-ME), and samples were left in the water bath for 10 mins. 2% β-ME was added to each sample and samples were boiled at 95°C for 3 min. The samples were run on a 6% SDS-PAGE gel, transferred to a membrane for 120 mins and blotted with anti-myc or anti-GluK2 antibodies.

#### Phos-tag gel electrophoresis

50mM Phos-tag gels (Fujifilm) were prepared following the manufacturer’s protocol. Cells were lysed in 200µl 2x sample buffer, sonicated and 20µl of each sample run on phos-tag gel. Gels were washed with 10mM EDTA in dH_2_O for 10 mins followed by Western blotting. Blots were blocked with milk and myc (Rb) antibody was used to detect GluK2 bands.

#### Quantitative Western Blots

To quantify the level of GluK2 SUMOylation, phosphorylation, and palmitoylation the ratio of SUMOylated to non-SUMOylated, phosphorylated to non-phosphorylated and palmitoylated to non-palmitoylated GluK2 was determined.

For palmitoylation state graphs, unconjugated, single and double palmitoylated bands were added, averaged and normalised to 100.

### Live Surface Labelling

Hippocampal neurons on 25 mm coverslips were transfected with YFP-Myc-GluK2 (WT or C2A) at DIV 9-10. At DIV 14-15, media was replaced with pre-warmed HBSS containing 2µM TTX (Tetrodotoxin) and 40µM GYKI53655 for 30 minutes to block neuronal activity and AMPARs, respectively. Coverslips were then incubated with 10µM kainate or vehicle control for 20 minutes. After kainate stimulation, neurons were incubated with chicken anti-GFP antibody (ab13970, 1:1000) in media for 20 minutes at 4°C. Next, neurons were washed quickly 5x in cold PBS before being fixed with pre-warmed (37°C) 4% PFA (ThermoFisher) for 20 minutes. Following fixation, neurons were washed 3x with PBS and then treated with glycine (100mM in PBS, Severn Biotech) for one minute after which they were washed 3x in PBS. Then, 3% BSA was used for blocking for 10 minutes before the incubation with Alexa 647-conjugated anti-chicken secondary antibody (1:400) in 3% BSA for one hour at room temperature in darkness. Neurons were then washed 3x with PBS and then permeabilised using 3% BSA and 0.1% Triton-X 100 (Fisher Scientific) for 10 minutes in darkness. After that, coverslips were incubated with chicken anti-GFP (1:1000) again to label the total transfected GluK2 for one hour in darkness. Neurons were washed with PBS (3x) before the addition of Cy2-conjugated anti-chicken secondary antibody (1:400, green) in darkness for one hour. Coverslips were then washed 3x before being mounted on prelabelled glass slides using Fluoromount-G with DAPI (eBioscience). Glass slides were kept in darkness to dry out for 24-48 hours before being imaged.

### Imaging Analysis

Confocal imaging was performed using a Leica SP5-AOBS confocal laser scanning microscope linked to a Leica DMI 6000 inverted epifluorescence microscope with the laser lines: 405 (blue for the nucleus), 488 (green for the total), 633 (far red for the surface). The transfected neurons were imaged by looking for the total labelling of GluK2 (green cells). A 63x oil immersion lens was used for image acquisition. Each image is composed of 5-6 stacks (0.4-0.5 µm stack interval) that were projected by maximum intensity. The untreated WT GluK2 condition was used to optimise the settings, which were kept constant throughout the same experiment. The immunofluorescence was quantified using ImageJ (FIJI).

### Statistical Analysis

All graphs and statistics were generated on GraphPad Prism version 9.0. The details of the statistical tests performed on each experiment are explained in the figure legend along with p-values and error bars.

## Acknowledgements

BPY is a Republic of Turkey Ministry of National Education funded PhD scholar. EMAM was a Hashemite University (Jordan) funded PhD scholar. We thank the BBSRC (BB/R00787X/), MRC (MR/W02036X/1), Wellcome Trust (220799/Z/20/Z) and Leverhulme Trust (RPG-2019-191) for financial support. We are grateful to the Wolfson Bioimaging Facility (University of Bristol).

## Author contributions

BPY performed all APEGS, phos-tag gel and biotinylation experiments. EMAM performed imaging and SUMOylation experiments. AJE initially proposed the link between palmitoylation and SUMOylation and provided creative input. RS helped with biochemistry. KAW made all lentiviral constructs. JMH and KAW supervised the study and wrote the paper and all authors participated in reading and editing the manuscript.

## References

1. A. Contractor, C. Mulle, G. T. Swanson, Kainate receptors coming of age: milestones of two decades of research. Trends Neurosci 34, 154–163 (2011).

2. J. Lerma, J. M. Marques, Kainate receptors in health and disease. Neuron 80, 292–311 (2013).

3. A. J. Evans, S. Gurung, J. M. Henley, Y. Nakamura, K. A. Wilkinson, Exciting Times: New Advances Towards Understanding the Regulation and Roles of Kainate Receptors. Neurochem Res 44, 572–584 (2019).

4. J. M. Henley, T. J. Craig, K. A. Wilkinson, Neuronal SUMOylation: mechanisms, physiology, and roles in neuronal dysfunction. Physiol Rev 94, 1249–1285 (2014).

5. C. Matute, Therapeutic Potential of Kainate Receptors. CNS Neurosci Ther 10.1111/j.1755-5949.2010.00204.x (2011).

6. B. Fritsch, J. Reis, M. Gasior, R. M. Kaminski, M. A. Rogawski, Role of GluK1 kainate receptors in seizures, epileptic discharges, and epileptogenesis. J Neurosci 34, 5765–5775 (2014).

7. G. Barthet et al., Presenilin and APP regulate synaptic kainate receptors. bioRxiv 10.1101/2022.02.03.478926, 2022.2002.2003.478926 (2022).

8. R. S. Petralia, Y. X. Wang, R. J. Wenthold, Histological and ultrastructural localization of the kainate receptor subunits, KA2 and GluR6/7, in the rat nervous system using selective antipeptide antibodies. J Comp Neurol 349, 85–110 (1994).

9. I. M. González-González, et al., Kainate Receptor Trafficking. WIRES Membrane Trasnsport and Signalling 1, 31–44 (2012).

10. M. Carta, S. Fievre, A. Gorlewicz, C. Mulle, Kainate receptors in the hippocampus. Eur J Neurosci 39, 1835–1844 (2014).

11. A. J. Evans, S. Gurung, J. M. Henley, Y. Nakamura, K. A. Wilkinson, Exciting Times: New Advances Towards Understanding the Regulation and Roles of Kainate Receptors. Neurochem Res 10.1007/s11064-017-2450-2 (2017).

12. Y. Nasu-Nishimura, H. Jaffe, J. T. Isaac, K. W. Roche, Differential regulation of kainate receptor trafficking by phosphorylation of distinct sites on GluR6. J Biol Chem 285, 2847–2856 (2010).

13. F. A. Konopacki et al., Agonist-induced PKC phosphorylation regulates GluK2 SUMOylation and kainate receptor endocytosis. Proc Natl Acad Sci U S A 108, 19772–19777 (2011).

14. S. E. Chamberlain et al., SUMOylation and phosphorylation of GluK2 regulate kainate receptor trafficking and synaptic plasticity. Nat Neurosci 15, 845–852 (2012).

15. G. D. Salinas et al., Actinfilin is a Cul3 substrate adaptor, linking GluR6 kainate receptor subunits to the ubiquitin-proteasome pathway. J Biol Chem 281, 40164–40173 (2006).

16. A. Maraschi et al., Parkin regulates kainate receptors by interacting with the GluK2 subunit. Nat Commun 5, 5182 (2014).

17. S. Martin, A. Nishimune, J. R. Mellor, J. M. Henley, SUMOylation regulates kainate-receptor-mediated synaptic transmission. Nature 447, 321–325 (2007).

18. D. S. Pickering, F. A. Taverna, M. W. Salter, D. R. Hampson, Palmitoylation of the GluR6 kainate receptor. Proc Natl Acad Sci U S A 92, 12090–12094. (1995).

19. B. A. Copits, G. T. Swanson, Kainate Receptor Post-Translational Modifications Differentially Regulate Association with 4.1N to Control Activity-Dependent Receptor Endocytosis. J Biol Chem M112.440719 [pii] 10.1074/jbc.M112.440719 (2013).

20. S. Martin, K. A. Wilkinson, A. Nishimune, J. M. Henley, Emerging extranuclear roles of protein SUMOylation in neuronal function and dysfunction. Nat Rev Neurosci 8, 948–959 (2007).

21. J. D. Nair, K. A. Wilkinson, J. M. Henley, J. R. Mellor, Kainate Receptors and Synaptic Plasticity. Neuropharmacology 10.1016/j.neuropharm.2021.108540, 108540 (2021).

22. A. J. Evans, S. Gurung, K. A. Wilkinson, D. J. Stephens, J. M. Henley, Assembly, Secretory Pathway Trafficking, and Surface Delivery of Kainate Receptors Is Regulated by Neuronal Activity. Cell Rep 19, 2613–2626 (2017).

23. J. Jin, X. Zhi, X. Wang, D. Meng, Protein palmitoylation and its pathophysiological relevance. J Cell Physiol 236, 3220–3233 (2021).

24. J. L. Daniotti, M. P. Pedro, J. Valdez Taubas, The role of S-acylation in protein trafficking. Traffic 18, 699–710 (2017).

25. L. Matt, K. Kim, D. Chowdhury, J. W. Hell, Role of Palmitoylation of Postsynaptic Proteins in Promoting Synaptic Plasticity. Front Mol Neurosci 12, 8 (2019).

26. Y. Fukata, M. Fukata, Protein palmitoylation in neuronal development and synaptic plasticity. Nat Rev Neurosci 11, 161–175 (2010).

27. I. Levental, M. Grzybek, K. Simons, Greasing their way: lipid modifications determine protein association with membrane rafts. Biochemistry 49, 6305–6316 (2010).

28. T. Kanadome, N. Yokoi, Y. Fukata, M. Fukata, Systematic Screening of Depalmitoylating Enzymes and Evaluation of Their Activities by the Acyl-PEGyl Exchange Gel-Shift (APEGS) Assay. Methods Mol Biol 2009, 83–98 (2019).

29. D. J. Speca, E. Diaz, Acyl-PEGyl Exchange Gel Shift Assay for Quantitative Determination of Palmitoylation of Brain Membrane Proteins. J Vis Exp 10.3791/61018 (2020).

30. K. A. Wilkinson, et al., “Using Lentiviral shRNA Delivery to Knock Down Proteins in Cultured Neurons and In Vivo” in Translational Research Methods in Neurodevelopmental Disorders, S. Martin, F. Laumonnier, Eds. (Springer US, New York, NY, 2022), 10.1007/978-1-0716-2569-9_1, pp. 1–17.

31. Z. Nagy, S. Comer, A. Smolenski, Analysis of Protein Phosphorylation Using Phos-Tag Gels. Curr Protoc Protein Sci 93, e64 (2018).

32. L. O’Donoghue, A. Smolenski, Analysis of protein phosphorylation using Phos-tag gels. J Proteomics 259, 104558 (2022).

33. E. I. Charych, L. X. Jiang, F. Lo, K. Sullivan, N. J. Brandon, Interplay of palmitoylation and phosphorylation in the trafficking and localization of phosphodiesterase 10A: implications for the treatment of schizophrenia. J Neurosci 30, 9027–9037 (2010).

34. A. Gauthier-Kemper, et al., Interplay between phosphorylation and palmitoylation mediates plasma membrane targeting and sorting of GAP43. Mol Biol Cell 25, 3284–3299 (2014).

35. P. Yan et al., Crosstalk of Synapsin1 palmitoylation and phosphorylation controls the dynamicity of synaptic vesicles in neurons. Cell Death Dis 13, 786 (2022).

36. B. A. Copits, G. T. Swanson, Kainate receptor post-translational modifications differentially regulate association with 4.1N to control activity-dependent receptor endocytosis. J Biol Chem 288, 8952–8965 (2013).

37. J. D. Nair, J. M. Henley, K. A. Wilkinson, Surface biotinylation of primary neurons to monitor changes in AMPA receptor surface expression in response to kainate receptor stimulation. STAR Protoc 2, 100992 (2021).

38. N. Yokoi et al., Identification of PSD-95 Depalmitoylating Enzymes. J Neurosci 36, 6431–6444 (2016).

39. A. Percher et al., Mass-tag labeling reveals site-specific and endogenous levels of protein S-fatty acylation. Proc Natl Acad Sci U S A 113, 4302–4307 (2016).

